# Activating an invertebrate bistable opsin with the all-trans 6.11 retinal analogue

**DOI:** 10.1101/2024.04.08.588240

**Authors:** Matthew J. Rodrigues, Oliver Tejero, Jonas Mühle, Filip K. Pamula, Ishita Das, Ching-Ju Tsai, Akihisa Terakita, Mordechai Sheves, Gebhard F.X. Schertler

## Abstract

Animal vision depends on opsins, a category of G protein-coupled receptor (GPCR) that achieves light sensitivity by covalent attachment to retinal. Typically binding as an inverse agonist in the 11-cis form, retinal photoisomerizes to the all-trans isomer and activates the receptor, initiating downstream signaling cascades. Retinal bound to bistable opsins isomerizes back to the 11-cis state after absorption of a second photon, inactivating the receptor. Bistable opsins are essential for invertebrate vision and non-visual light perception across the animal kingdom. While crystal structures are available for bistable opsins in the inactive state, it has proven difficult to form homogeneous populations of activated bistable opsins either via illumination or reconstitution with all-trans retinal. Here we show that a non-natural retinal analogue, all-trans retinal 6.11 (ATR6.11), can be reconstituted with the invertebrate bistable opsin, Jumping Spider Rhodopsin-1 (JSR1). Biochemical activity assays demonstrate that ATR6.11 functions as an agonist of JSR1. ATR6.11 binding also enables complex formation between JSR1 and downstream signaling partners. Our findings demonstrate the utility of retinal analogues for biophysical characterization of bistable opsins, which will deepen our understanding of light perception in animals.

## Introduction

Jumping Spider Rhodopsin-1 is a light-sensitive GPCR that the jumping spider requires for depth perception (1). JSR1 achieves light sensitivity by covalent binding of 11-cis retinal chromophore to a lysine side chain via a protonated Schiff base (PSB) (2). Photoisomerization of retinal to the all-trans isomer, triggers conformational rearrangements in the receptor, resulting in adoption of an active conformation capable of catalyzing nucleotide exchange in intracellular G proteins (2, 3). The receptor thereby transduces an optical signal to initiate cellular signaling cascades. Unlike vertebrate visual opsins, which are bleached and lose the retinal after photo-activation, all-trans retinal bound to invertebrate opsins and non-visual vertebrate opsins reverts to the 11-cis state after a second photo-isomerisation event (4). They are therefore classified as ‘bistable’ due to the thermal stability of the PSB in both the inactive and active states. Bistable opsins are potential optogenetic switches to control G protein signaling pathways, and *in vivo* studies have demonstrated the capability of JSR1 as an optogenetic tool to manipulate neuronal signaling in animals (5).

High-resolution crystal structures of JSR1 and squid opsin have provided insights into the architecture of the retinal binding site in bistable opsins (3, 6). These structures identify several structural differences between monostable (bleachable) and bistable (non-bleachable) opsins, particularly in the retinal binding site (3, 6). In particular, the two categories of receptor have evolved distinct counterion systems that stabilize the positive charge of the PSB within the hydrophobic core of the receptor. There are currently no active state structures of bistable opsins interacting with downstream signaling partners and it is therefore unclear how these receptors change conformation after retinal isomerization. Structural studies depend on obtaining a homogeneous population of the receptor in the active state. In the case of JSR1, it has not been possible to reconstitute the receptor with the native agonist, all-trans retinal. In addition, due to the overlapped absorption maxima (λ_max_) of the inactive and active states, illumination of the receptor generates a mixture of states not easily amenable to structural characterization. It is not uncommon for the λ_max_ of bistable opsins to be similar in the active and inactive states, as shown for JSR1, melanopsin and squid rhodopsin (2, 7, 8). The present study focuses on the use of locked retinal analogues to activate JSR1 without light.

## Results

JSR1 was expressed recombinantly in the apo state, and addition of 9-cis retinal (Figure 1a) to the cell lysate enabled reconstitution with the inverse agonist, as evidenced by the 505 nm absorption maximum (Figure 1b). Adding the endogenous agonist, all-trans retinal, to cell lysate containing JSR1 failed to reconstitute the protein with the chromophore and to form the protein-chromophore covalent bond (Figure 1b). We therefore sought for non-natural agonists of JSR1 that could activate the receptor. We attempted reconstitution of JSR1 with 9-cis retinal 6.11 (9CR6.11) and ATR6.11, which have a six-membered ring cyclized around the 11-double bond of retinal (Figure 1a). JSR1 bound both ATR6.11 and 9CR6.11, as demonstrated by the absorption spectrum of the protein after purification (Figure 1b). The 513 nm and 509 nm λ_max_ of JSR1 bound to ATR6.11 and 9CR6.11 respectively suggests that they are covalently bound to JSR1 via a PSB (Figure 1b).

**Figure 1.**
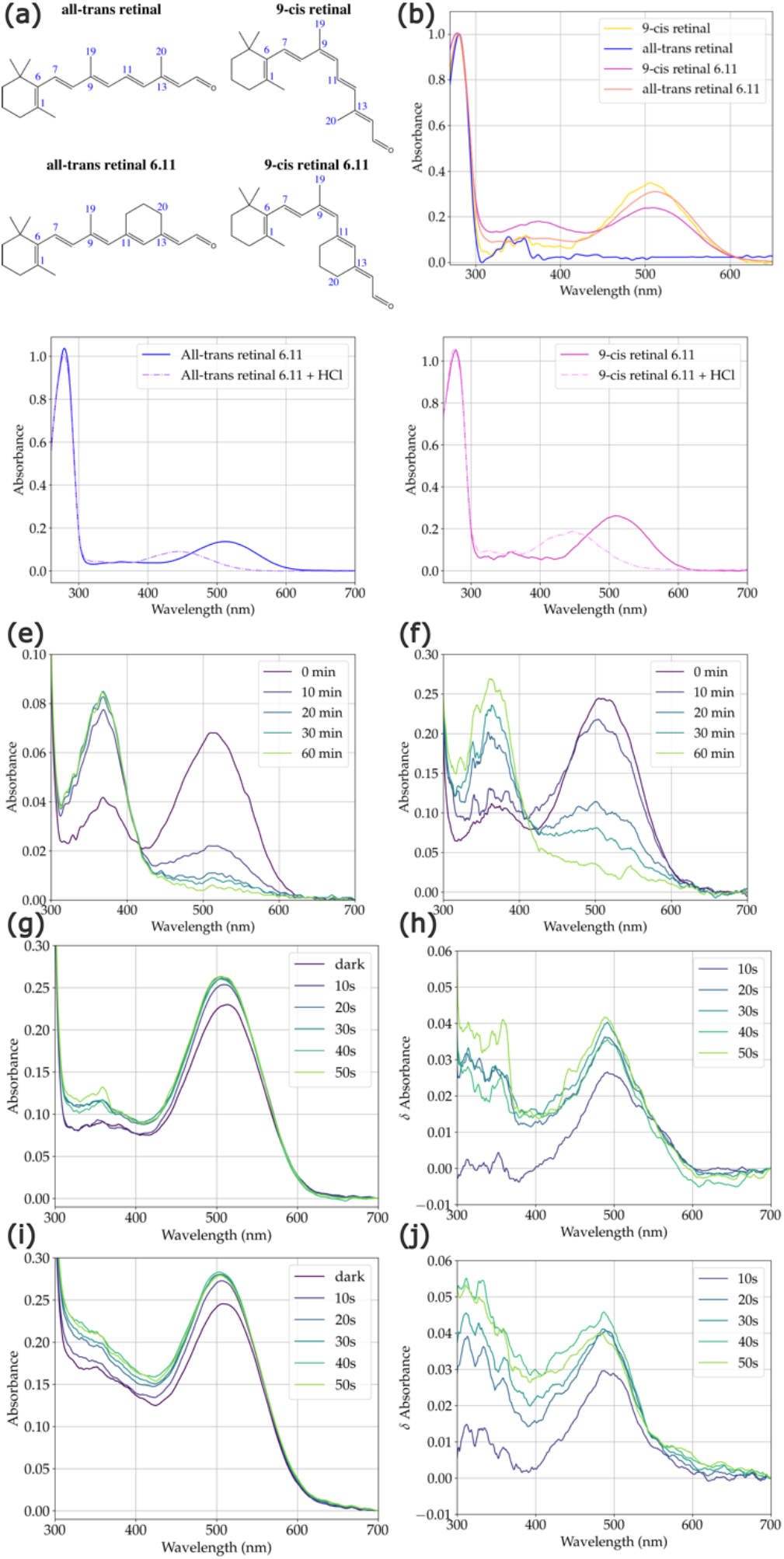
(a) Chemical structures of all-trans retinal, 9-cis retinal, all-trans retinal 6.11 and 9-cis retinal 6.11. (b) UV-Vis spectra of purified JSR1 after reconstitution with 9-cis retinal, all-trans retinal, 9-cis retinal 6.11 and all-trans retinal 6.11. (c) UV-Vis spectra of JSR1/ATR6.11 before and after acid denaturation. (d) UV-Vis spectra of JSR1/9CR6.11 before and after acid denaturation. (e) UV-Vis spectra of JSR1/ATR6.11 after addition of hydroxylamine. (f) UV-Vis spectra of JSR1/9CR6.11 after addition of hydroxylamine. (g, h) UV-Vis spectra and difference spectra of JSR1/ATR6.11 after repeated illumination at 519 nm. (i, j) UV-Vis spectra and difference spectra of JSR1/9CR6.11 after repeated illumination at 519 nm.

We performed an acid denaturation experiment to confirm that ATR6.11 and 9CR6.11 were covalently bound to JSR1 via a protonated Schiff base. The λ_max_ shifts from ≈510 nm to 445 nm (Figure 1c & 1d) for both retinal isomers after addition of HCl, as expected if the protein unfolds and the PSB becomes solvent exposed (8). The molar extinction coefficients of JSR1/ATR6.11 and JSR1/9CR6.11 were determined experimentally using a hydroxylamine assay. Addition of hydroxylamine to the opsin cleaves the PSB and releases the retinal oxime (λ_max_ 365 nm). The extinction coefficient is calculated using the extinction coefficients determined for ATR6.11 and 9CR6.11, and the ratio between the decrease in absorbance at 509 nm and increase at 365 nm. We determined the extinction coefficient of JSR1/ATR6.11 to be 49,500 M^−1^ cm^−1^ at 509 nm (Figure 1e), while JSR1 in complex with 9CR6.11, has an extinction coefficient of 37,430 M^−1^cm^−1^ at 509 nm (Figure 1f). The higher extinction coefficient of ATR6.11 compared to 9CR6.11 corresponds well to the higher extinction coefficient of JSR1 bound to all-trans retinal compared to 9-cis, 37,560 M^−1^ cm^−1^ and 32,660 M^−1^ cm^-1^ respectively (2).

While the C_11_=C_12_ bond of the retinal analogues is constrained and cannot isomerize, other bonds may still be susceptible to photo-isomerization. We measured UV-Vis spectra of JSR1 bound to the retinal analogues before and after illumination at 519 nm for defined time periods. In the case of JSR1/ATR6.11, we observe a marked increase in absorption within the first 10 seconds of illumination (Figure 1g), with the greatest increase in absorption at 491 nm (Figure 1h). We also observe increased absorption by JSR1/9CR6.11 within the first 10 seconds of illumination (Figure 1i), with the formation of a photoproduct with a higher extinction coefficient at 488 nm (Figure 1j). As the 9-cis bond is liable to isomerize, it is possible that an all-trans isomer is formed, although the formation of a di-cis species cannot be excluded (9). While the molecular bases of these changes have not been resolved, it is evident that both retinal 6.11 analogues remain light sensitive after binding to JSR1.

After confirming covalent binding of both retinal analogues, we sought to determine whether they function as agonists. Based on phylogenetic analysis, JSR1 is expected to catalyze exchange of GDP for GTP in the Gq heterotrimer when in the active state (10), although it also couples to human Gi *in vitro* (3). A GTPase-Glo assay was used to measure the amount of GTP remaining after incubating GTP with JSR1 and either Gq or Gi heterotrimer, and hence the ability of JSR1 to catalyze nucleotide exchange in the G proteins. In the absence of JSR1, Gi demonstrates a markedly higher intrinsic rate of nucleotide turnover than Gq, with almost 60% of nucleotides depleted compared to 15% for Gq (Figure 2). Addition of JSR1/9-cis retinal causes no increase in the rate of nucleotide exchange in the Gi or Gq heterotrimers (Figure 2a), as expected, since 9-cis retinal stabilizes JSR1 in an inactive conformation (3). Steady state illumination of 9-cis retinal bound JSR1 generates a dynamic equilibrium of JSR1 conformational states, due to the overlapped absorption profiles of the 9-cis, 11-cis and all-trans isomers, with up to 73% of the molecules adopting the active all-trans conformation (2). In keeping with this, we observe an increase in GTP depletion by Gi from 57% in the presence of JSR1/9-cis retinal to 92% after illumination (Figure 2a). Similarly, nucleotide depletion by Gq increases from 7% to 44%, a clear marker of light-induced receptor activation (Figure 2a).

**Figure 2.**
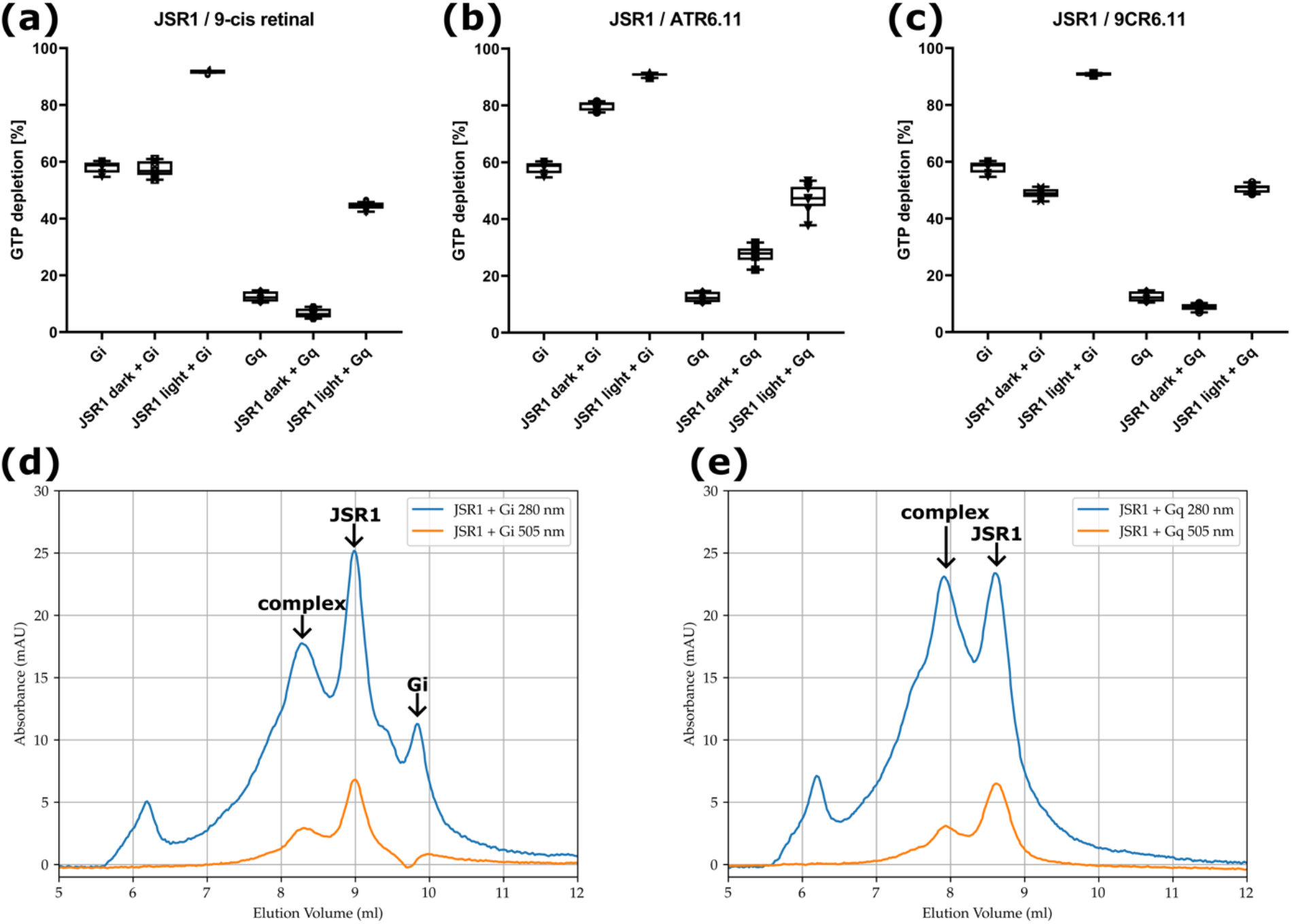
GTP depletion by the human Gi and Gq heterotrimers in the presence and absence of JSR1 bound to (a) 9-cis retinal, (b) all-trans retinal 6.11, (c) 9-cis retinal 6.11, with and without illumination. Size exclusion chromatography (SEC) profiles for dark state JSR1/ATR6.11 incubated with the (d) human Gi heterotrimer and (e) human Gq heterotrimer.

In contrast, JSR1/ATR6.11 catalyzes nucleotide exchange without illumination, increasing GTP depletion in the presence of Gi from 58% to 80% and from 13% to 28% in the presence of Gq (Figure 2b). Illumination of JSR1/ATR6.11 causes a further increase in activity in Gi and Gq to 91% and 47% respectively (Figure 2b). While the C_11_=C_12_ double bond of the ligand is constrained in a trans configuration, absorption of light may induce structural changes in the chromophore that increase its potency as an agonist of JSR1. In the dark state, JSR1/9CR6.11 demonstrates no catalytic effect on nucleotide depletion by Gi or Gq (Figure 2c), similar to the effect of 9-cis retinal. Illumination of the sample allows the receptor to catalyze nucleotide exchange in Gi and Gq to similar levels observed for illuminated JSR1/9-cis retinal and JSR1/ATR6.11, with 91% depletion in the presence of Gi and 51% with Gq (Figure 2c). The light induced increase in nucleotide depletion may be caused by isomerization around the C_9_=C_10_ double bond of 9CR6.11, yielding ATR6.11 chromophore. However, light induced formation of di-cis retinal analogues has also been shown to activate monostable opsins previously (9).

Finally, we sought to determine whether JSR1/ATR6.11 would form a stable complex with the human Gi and Gq heterotrimers. Purified receptor and G protein was incubated together in the presence of apyrase. The receptor and the G protein heterotrimers were subjected to size exclusion chromatography (SEC) after incubation together. Both samples show a pronounced peak at an elution volume of ∼8.8 ml corresponding to the receptor alone, with a relatively high ratio of absorbance at 505 nm compared to the 280 nm (Figure 2d & 2e). An additional peak is visible at 7.8 ml at both wavelengths when JSR1/ATR6.11 is incubated with the Gq heterotrimer (Figure 2e). The shorter retention volume indicates formation of a higher molecular weight complex. Similarly, when JSR1/ATR6.11 was incubated with the Gi heterotrimer, a species with a shorter retention volume (8.3 ml) formed with absorbance at both 280 nm and 505 nm (Figure 2d). SDS-PAGE analysis confirmed formation of the full complex of JSR1/ATR6.11 with the Gi and Gq heterotrimers. This demonstrates that ATR6.11 is a sufficiently potent agonist to induce complex formation with downstream signaling partners. We note that significant populations of the receptor and G protein remain unbound to each other in the Gi sample and this complex may therefore be less stable than the Gq complex (Figure 2d).

## Discussion

Locked retinal analogues have been used to investigate the photochemistry underlying signaling by retinal proteins, both with microbial opsins (11, 12) and monostable vertebrate opsins (9, 13–15). We show that their use can be expanded to biochemical and biophysical studies of bistable opsins. Despite their physiological importance, our understanding of the mechanisms by which these proteins achieve bistability remains limited, largely due to difficulties with recombinant expression and successful reconstitution of the receptors with their native chromophores. We demonstrate that retinal analogues function as tool molecules to inform our understanding of bistable opsins. In particular, we show that reconstitution of ATR6.11 with JSR1 forms a population of activated receptor suitable for structural and biophysical studies. The underlying reason for the selective binding of ATR6.11 over all-trans retinal remains unclear. ATR6.11 may simply bind with higher affinity, or the aldehyde group may be better positioned to form a Schiff base bond with the binding site lysine residue. It should be noted that most bistable opsins have not evolved to bind all-trans retinal and light-independent binding of a natural agonist would contribute to dark noise, with light independent receptor activation. We envisage that these compounds will be of use to the broader opsin community, as their ability to activate a given protein can be easily tested using biochemical and cellular signaling assays.

## Materials and Methods

Full materials and methods may be found in the Supplementary Information.

## Acknowledgments

We would like to thank Mara Wieser, Dr. Elena Lesca and Dr. Niranjan Varma for their contributions in JSR1 biochemistry. This project has received funding from the European Research Council (ERC) under the European Union’s Horizon 2020 research and innovation programme (Grant agreement No. 951644 to GFXS, and Marie Skłodowska-Curie grant agreement No. 701647 to MJR). M.S. thanks the Kimmelman Center for Biomolecular Structure and Assembly for partial support. M.S. holds the Katzir-Makineni Chair in Chemistry.

