## Supplementary Information for "Activating an invertebrate bistable opsin with the all-trans 6.11 retinal analogue"

**Matthew J. Rodrigues *et al.***

**This supplement includes:**

Extended Materials and Methods

### Extended Materials and Methods

*Synthesis of retinal analogues.* All-trans retinal 6.11 was synthesized as described previously (1). 9-cis retinal 6.11 was prepared similarly but from the cis isomer of the appropriate ketone ((Z)-3-[E-2-Methyl-4-(2,6,6-trimethyl-1-cyclohexenyl)-3-butenel-diene]-2-cyclohexene-1-one) as previously described (1).

*JSR1 expression and purification.* Wild-type Jumping Spider Rhodopsin isoform-1 (JSR1) from *Hasarius adansoni* tagged with a C-terminal 1D4 epitope (2), was recombinantly expressed in HEK293 GnTi− cells as described previously (3). Cells were harvested by centrifugation at 500 g and the pellets were stored at -80 °C. The cells were later thawed in lysis buffer (50 mM HEPES pH 6.5, 150 mM sodium chloride, 3 mM magnesium chloride) and mechanically disrupted in a Dounce homogeniser. All following steps were performed under dim red light conditions. Retinal was added to the lysate (9-cis retinal 30 μM (98% grade; Sigma-Aldrich, MA, USA), 9-cis retinal 6.11 10 μM, all-trans retinal 6.11 10 μM) and the suspension was incubated at 4 °C overnight. Dodecyl-β -D-maltoside (DDM) powder (DDM Solgrade; Anatrace, OH, USA) was then added to a final concentration of 1% (w/v) to solubilise the receptor and centrifuged at 100,000 g for one hour. The supernatant was incubated with CNBr-Activated Sepharose resin (GE Healthcare Life Science, Germany) coupled to anti-1D4 antibody (Cell Essentials, MA, USA) at 4 °C overnight, the resin was then loaded into a glass Econo-Column (Bio-Rad Laboratories, CA), and washed with 30 column volumes of wash buffer (50 mM HEPES pH 6.5, 150 mM sodium chloride, 3 mM magnesium chloride, 0.01% DDM (w/v)) before the addition of 1.5 column volumes of elution buffer (50 mM HEPES pH 6.5, 150 mM sodium chloride, 3 mM magnesium chloride, 0.01% DDM (w/v), 800 μM 1D4 peptide (Peptide 2.0, VA, USA)) to the resin. The slurry was incubated at 4 °C overnight and the protein was eluted from the column the following day. Residual protein was eluted from the resin after the addition of three more column volumes of wash buffer. All eluted fractions were pooled.

*UV-Vis spectroscopy*. UV-Vis spectra of 1D4-purified JSR1 samples were measured using a Shimadzu UV-2401PC spectrophotometer with 1D4 purified JSR1. The spectra were smoothed using a Savitzky-Golay function (4) with a 21 nm window and second order polynomial within SciPy (5), and normalised to zero absorbance at the wavelength with the lowest absorption reading and plotted using Matplotlib (6) in a custom Python script.

*Acid denaturation assay.* UV-Vis spectra were measured of purified JSR1 reconstituted with either 9-cis or all-trans retinal 6.11, in a volume of 70 μl. Hydrochloric acid was added to the protein samples to a final concentration of 40 mM and total volume of 100 μl. A second UV-Vis spectrum was measured after the addition of the acid. Spectra were smoothed using a Savitzky-Golay function (4) with a 21 nm window and second order polynomial. The baseline absorbance at 700 nm was subtracted, and the smoothed spectra were scaled to an absorbance of 1.0 at 280 nm within SciPy (5), and plotted using Matplotlib (6) in a custom Python script.

*Extinction coefficient of retinal oximes*. A serial dilution of the two retinal analogues was performed after an initial 70-fold dilution in wash buffer, to accurately determine the stock concentration of the compounds. A straight line was fitted to all values to determine the stock concentration using the extinction coefficients (ε), (9-cis retinal 6.11 ε = 27,600$^{-1}$ cm$^{-1}$, all-trans retinal 6.11 ε = 27,700 M$^{-1}$ cm$^{-1}$), which were determined by measuring the absorption of light by a weighed quantity of the retinal analogues. The extinction coefficients of the all-trans retinal 6.11 and 9-cis retinal 6.11 oximes were then determined by diluting the retinal analogues by a factor of 70 in a hydroxylamine assay buffer composed of 50 mM HEPES pH 6.5, 150 mM sodium chloride, 3 mM magnesium chloride, 0.01% DDM (w/v) and 1 M hydroxylamine. The mixture was incubated at 4 °C for 20 minutes to allow the reaction to occur. A UV-Vis spectrum was then measured from each retinal oxime sample. A serial dilution with 2x dilution factor was performed and the spectra were measured at each concentration until the absorption at 380 nm was negligible. The absorption at 380 nm was plotted as a function of retinal concentration and the gradient of a straight line fitted to the data was used to calculate the extinction coefficient of each retinal oxime at 380 nm.

*Hydroxylamine assay*. For each retinal isomer, 30 µl hydroxylamine assay buffer was added to 70 µl of purified JSR1 reconstituted with retinal, UV-Vis spectra were measured at five-minute intervals for one hour. The change in absorbance at 365 nm was plotted against the change in absorbance 509 nm and the ratio was used to determine the extinction coefficient of each retinal isomer while bound to JSR1.

*UV-Vis spectroscopy of illuminated JSR1/retinal 6.11.* UV-Vis spectra were measured of JSR1/all-trans retinal 6.11 and JSR1/9-cis retinal 6.11 in the dark state. Protein samples were illuminated at 519 nm with a mean irradiance of 1.00 mW $cm^{-2}$ using an optoWELL LED light source (Opto Biolabs GmbH, Freiburg im Breisgau, Germany) for 10 second intervals. A UV-Vis spectrum was measured after each illumination, spectra were smoothed using a Savitzky-Golay function (4) with a 21 nm window and second order polynomial. The baseline absorbance at 700 nm was subtracted, and the smoothed spectra were scaled to an absorbance of 1.0 at 280 nm within SciPy (5) in a custom Python script. Difference spectra were calculated by subtracting the smoothed and normalised dark state spectrum from the smoothed and normalised spectra measured from the illuminated samples. Spectra and difference spectra were plotted using Matplotlib (6).

*$G\alpha i_1\beta_1\gamma_1$ expression and purification.* Human Gαi subunit ($G\alpha i_1$) with an N-terminal TEV protease-cleavable deca-histidine tag was expressed in *E. coli* BL21 (DE3) cells (Sigma-Aldrich: CMC0014) and purified as described previously (7). The $G\beta_1\gamma_1$ subunits were separated from the transducin G protein heterotrimer, which was purified from bovine retinae as described by Maeda *et al* (8). Human $G\alpha i_1$ and bovine $G\beta_1\gamma_1$ were incubated together to generate the $G\alpha i_1\beta_1\gamma_1$ heterotrimer used for complex formation with JSR1.

*Gq expression.* Genes encoding the wild type human Gαq subunit and human GST-Ric8A were cloned into a pAC8REDNK vector for insect cell expression. Separately, genes encoding human Gβ1 modified with an N-terminal 10xHis tag followed by a HRV 3C cleavage sequence and the unmodified human $G\gamma_2$ were cloned into a separate pAC8REDNK vector. Baculoviruses were produced in Sf9 insect cells using the FlashBac technology (Oxford Expression Technologies, Oxford, UK). 1.0 x $10^6$ Sf9 cells were co-transfected with 1.5 µg plasmid DNA and 500 ng linearized Bac10:KO1629 viral DNA using Cellfectin II (Invitrogen, MA, USA) following the manufacturer's protocol. Cells were incubated for five hours at 27 °C with the transfection mixture before it was replaced by fresh SF900II SFM insect cell medium (Invitrogen, MA, USA) supplemented with 1% Pen/Strep (PAN Biotech, Aidenbach, Germany). After five days of incubation at 27 °C, V0 generation virus was harvested by centrifugation at 800 x g and 20 °C, the supernatant supplemented with 1% Pen/Strep and 10% FCS (Sigma-Aldrich, MA, USA) and stored in the dark at 4 °C. The viruses were first amplified by infecting new Sf9 cultures at 1.0 x $10^6$ cells/ml in SF900II SFM medium with 10% V0 generation viruses. The V1 generation viruses were harvested and stored 72 hours post-infection as described above for the V0 viruses, however, only 1% FCS was used as a supplement. A second round of virus amplification was performed by infecting new Sf9 cultures at 1.0 x $10^6$ cells/ml in SF900II SFM medium with 1% V1 generation viruses. The V2 generation viruses were harvested and stored as described for the V1 viruses. The Gq heterotrimer was expressed in High Five insect cells (Invitrogen, MA, USA) by co-infection with the virus encoding the Gαq subunit and human GST-Ric8A and the virus encoding for N-terminally 10x-His tagged human $G\beta_1$ and the unmodified human $G\gamma_2$. Large-scale expression was conducted at a cell density of 3.0 x $10^6$ cells/ml in SF900II SFM medium using 2 L Erlenmeyer flasks. The viruses were added in an optimal ratio and the cells were cultured at 27 °C and 120 rpm and harvested 48 hours post-infection by centrifugation. Pellets were flash frozen in liquid nitrogen and stored at -80 °C.

*Gq purification.* The frozen cell pellet was thawed and resuspended in a buffer containing 20 mM HEPES pH 7.5, 100 mM NaCl, 3 mM $MgCl_2$, 100 µM EDTA, 5 mM β-mercaptoethanol and 20 µM GDP supplemented with DNase I (10 µg/ml) and cOmplete protease inhibitor cocktail tablets (Roche, Basel, Switzerland). Cells were mechanically disrupted using a Microfluidizer (IDEX Health & Science, NY, USA) and subsequently centrifuged at 185,000 rcf for 45 min. The pellet was resuspended in 20 mM HEPES, 100 mM sodium chloride, 20 mM imidazole, 3 mM magnesium chloride, 5 mM β-mercaptoethanol and 20 µM GDP supplemented with cOmplete protease inhibitor cocktail tablets and solubilized by adding 1% sodium cholate (Sigma-Aldrich) and 0.05% DDM (Anatrace, OH, USA). After incubation at 4 °C for 1 h, insoluble material was removed by centrifugation at 185,000 rcf for 45 min. The clarified supernatant was incubated with 5 ml of TALON resin (Takara Bio, CA, USA) for 90 min at 4 °C. The resin was collected in a glass Econo-Column (Bio-Rad Laboratories, CA, USA) and the detergent was gradually exchanged to 0.1 % DDM. The resin was washed with 10 column volumes of IMAC buffer A (20 mM HEPES pH 7.5, 100 mM sodium chloride, 20 mM imidazole, 1 mM magnesium chloride, 5 mM β-mercaptoethanol, 20 µM GDP and 0.01 % DDM) and 10 column volumes of IMAC buffer B (IMAC buffer A with 300 mM sodium chloride). The sample was eluted with IMAC buffer A supplemented with 300 mM Imidazole. His-tagged HRV 3C protease was added (1:50 w/w; HRV 3C protease : Gq heterotrimer) to the eluent and dialyzed against dialysis buffer (20 mM HEPES pH 7.5, 100 mM sodium chloride, 1 mM magnesium chloride, 100 µM TCEP, 20 µM GDP and 0.05 % DDM) overnight at 4 °C. Dialysate was collected and supplemented with imidazole to a final concentration of 20 mM before incubation with 5 ml TALON resin for 1 h at 4 °C. The resin was transferred to a gravity flow column and the flow through was collected. The resin was then washed with one column volume of dialysis buffer supplemented with 20 mM imidazole. The flow through and wash were combined and loaded to a HiTrap Q FF 1 ml column (Cytiva, MA, USA). The column was washed with 15 column volumes of Q buffer A (20 mM HEPES pH 7.5, 100 mM sodium chloride, 1 mM magnesium chloride, 100 µM TCEP, 20 µM GDP and 0.05 % DDM) and the sample was eluted with a linear gradient over 30 column volumes with Q buffer B (Q bufferA with 1000 mM sodium chloride). The relevant fractions were pooled and concentrated using a 50 kDa molecular weight cut-off (MWCO) Amicon Ultra Centrifugal Filter (Millipore, MA, USA). The concentrated sample was injected into a SRT-C 300 SEC-300 10x300 mm column (Sepax Technologies, DE, USA) pre-equilibrated with SEC buffer (20 mM HEPES pH 7.5, 100 mM sodium chloride, 1 mM magnesium chloride, 100 µM TCEP, 20 µM GDP and 0.02% DDM). Fractions containing the Gq heterotrimer were pooled and concentrated using a 50 kDa MWCO Amicon Ultra Centrifugal Filter (Millipore, MA, USA) before adding 10% glycerol (v/v). The sample was flash frozen in liquid nitrogen and stored at -80 °C.

*GTP Glo Assay.* GTPase Glo Assay reactions (Promega, WI, USA) (9), were set up in 384-well, low-volume, PS, white, solid-bottom plates (Greiner Bio-One, Kremsmünster, Austria) and carried out at 20 °C under dim-red light conditions. Each condition was prepared in octuplicates. 2.5 µl of protein solution were mixed with 2.5 µl of 2 µM GTP solution. Reference samples contained either 2 µM G protein or 0.2 µM JSR1 with either 9-cis retinal, all-trans retinal 6.11 or 9-cis retinal 6.11 (dark and light activated). Light samples were illuminated at a wavelength of 519 nm for 10 min using an Optowell device (Opto Biolabs GmbH, Freiburg im Breisgau, Germany). Test samples contained 2 µM G protein and 0.2 µM JSR1 with the respective ligand (dark and light activated). A control was performed with no protein. The plate was sealed and incubated for 120 minutes shaking at 500 rpm. Subsequently, 5 µl of GTP-Glo reagent was added to each well and the reaction was carried out for 30 minutes. Finally, 10 µl of Detection Reagent was added to each well. Luminescence was measured after 10 minutes in a PheraStar FSX (BMG Labtech, Ortenberg, Germany) plate reader (Optical module: LUM plus, Gain: 3600, focal height: 14.5, 1 second/well). Outliers were identified using the Grubbs test (10). GTP hydrolysis was calculated in relation to the control sample.

*Complex Formation and Size Exclusion Chromatography.* JSR1 bound to all-trans retinal 6.11 was purified as described above, however after washing the 1D4 column with wash buffer it was washed with a further 15 column volumes of LMNG (lauryl maltose neopentyl glycol) wash buffer (50 mM HEPES pH 6.5, 150 mM sodium chloride, 3 mM magnesium chloride, 0.0015% LMNG (w/v)). The receptor was then eluted from the 1D4 column in an elution buffer containing LMNG (50 mM HEPES pH 6.5, 150 mM sodium chloride, 3 mM magnesium chloride, 0.0015% LMNG, 800 µM 1D4 peptide). The 1D4 purified receptor was concentrated in a 15 ml 50 kDa MWCO centrifugal concentrator (Merck Millipore, MA, USA) to a concentration of 8 µM and incubated with equimolar amounts of either preformed $G\alpha i_1\beta_1\gamma_1$ or preformed $G\alpha q\beta_1\gamma_2$ heterotrimer on ice under dim red-light conditions for two hours in the presence of apyrase enzyme (25 mU/ml) to degrade GTP and GDP present in the solution. The mixture was concentrated to a volume of 180 µl in a 0.5 ml 100 kDa MWCO centrifugal concentrator (Merck Millipore, MA, USA). The sample was injected onto a 24 ml Superdex 200 Increase 10/300 GL size exclusion chromatography (SEC) column pre-equilibrated in SEC buffer (20 mM HEPES pH 7.5, 100 mM sodium chloride), and run at 0.35 ml/min under ambient light conditions.
